# The complete mitochondrial genome of *Hanabira yukibana* Lau, Stokvis, Imahara & Reimer, 2019 (Cnidaria: Anthozoa: Octocorallia)

**DOI:** 10.1101/2025.09.27.678955

**Authors:** Yuki Yoshioka, Tatsuki Koido, Megumi Kanai, Shogo Gishitomi, Natsuki Watanabe, Noriyuki Satoh, Tomofumi Nagata

**Affiliations:** Marine Genomics Unit, Okinawa Institute of Science and Technology Graduate University, Onna, Okinawa 904-0495, Japan; Kuroshio Biological Research Foundation, Otsuki, Kochi 788-0333, Japan; Incorporated Foundation Okinawa Environment Science Center, Urasoe, Okinawa 901-2111, Japan

**Keywords:** Octocorals, Malacalcyonacea, Okinawa Island, molecular phylogenetics

## Abstract

In this study, we sequenced the complete mitochondrial genome (mitogenome) of *Hanabira yukibana* (family Clavulariidae), collected in Okinawa, Japan. The complete mitochondrial genome of this specimen was 18,690 bp in length and contained 17 genes (14 protein-coding genes, two rRNA genes, and one tRNA gene). It exhibited gene order pattern A, which is the most common arrangement among octocorals. Molecular phylogenetic analysis indicated that *H. yukibana* is clustered with the genus *Clavularia*, including *Clavularia inflata* (the family Clavulariidae), consistent with the previous research using partial mitochondrial genes. Our study provides an additional mitochondrial genomic resource that contributes to comparative studies of octocoral mitogenomes.

## Introduction

The anthozoan class Octocorallia Haeckel, 1866 comprises more than 3,500 described species (Williams and Cairns, 2019) inhabiting a wide range of marine environments, from shallow coral reefs to the deep sea (Cairns, 2007; Dinesen, 1983). The Octocorallia is phylogenetically divided into two clades, the orders Malacalcyonacea and Scleralcyonacea (formerly known as Alcyonacea, Pennatulacea, or Helioporacea) (McFadden et al., 2022). The genetic feature unique to octocorals is the presence of the *mutS* gene in their mitogenome (*mt*-*mutS*), a homolog of the epsilonproteobacterial mismatch repair gene *mutS* (Bilewitch and Degnan, 2011; McFadden et al., 2010; Pont-Kingdon et al., 1995).

*Hanabira yukibana* Lau, Stokvis, Imahara & Reimer, 2019, a member of the family Clavulariidae Hickson, 1894, was first described from Okinawa, Japan (Lau et al., 2019). Similar to many shallow-reef octocorals, this species harbors symbiotic algae (Lau et al., 2019). The genus name *Hanabira* (“petal” in Japanese) refers to the petal-like shape of the polyp tentacles, whereas the species name *yukibana* (“snow flower” in Japanese) alludes to the delicate sheen of the polyps. Because species identification in octocorals based solely on morphology remains challenging, genomic resources are indispensable for accurate taxonomic resolution. To date, mitogenome for this species have not been reported. In the present study, we report the complete mitogenome of *H. yukibana* and perform molecular phylogenetic analyses.

## Materials and Methods

We collected a colony of *H. yukibana* in the reef slope (at 14.3 m in depth) near Yamakawa Port (latitude: 26.679029 and longitude: 127.880824), Okinawa-jima, Japan on 7 June 2024. A specimen was deposited at Kuroshio Biological Research foundation, Kochi, Japan (KBF) in the octocoral collection (OA) (https://kuroshio.or.jp/, contact parson: Tatsuki Koido) under the voucher number “KBF-OA-00429”. Sample was preserved with 99.5% ethanol immediately after collection. Species was identified based on colony and polyp morphology, and observation of sclerites using scanning electron microscopy (JCM-7000 NeoScope™, JEOL Ltd.). Genomic DNA was extracted from the anthocodial using a Maxwell RSC Blood DNA Kit (Promega). Sequence libraries were constructed with NEBNext Ultra II FS DNA PCR-free Library Prep Kit for Illumina (New England Biolabs) according to the manufacturer’s protocol and were sequenced on an Illumina NovaSeq X, with 150-bp paired-end reads. Illumina sequence adaptors and low-quality sequences (quality cutoff=20) were trimmed with CUTADAPT v4.3 (Martin, 2011). Cleaned reads were assembled with GetOrganelle v1.7.7.0 (Jin et al., 2020). The sequencing depth was calculated with BamDeal v0.27 (https://github.com/BGI-shenzhen/BamDeal). Mitochondrial gene annotation was performed with MITOS2 (Bernt et al., 2013) and missing annotations were manually curated. Complete mitogenome with gene annotation were visualized with OrganelleGenomeDRAW (Greiner et al. 2019). We performed molecular phylogenetic analysis following Yoshioka et al. (2025). The families Ideogorgiidae and Sarcodictyonidae belonging to Scleralcyonacea were used for outgroups.

## Results

We successfully obtained the complete mitogenome of *H. yukibana* with an average coverage of 279x (Figure 2). The mitogenome was 18,690 bp and encoded 14 protein-coding genes (*nad1*–*6, nad4l, cox1*–*3, atp6, atp8, cob*, and *mt-mutS*), two rRNA genes (*rrnS* and *rrnL*), and one tRNA gene (*trnM*) (Figure 2). To date, 14 gene order rearrangements, recognized as patterns A to M and F1, have been discovered in Octocorallia (Brockman and McFadden, 2012; Brugler and France, 2008; Hogan et al., 2019; Pante et al., 2013; Park et al., 2012; Poliseno et al., 2025; Uda et al., 2011; Yoshioka et al., 2025). The mitogenome of *H. yukibana* exhibited gene order pattern A, which is the most common arrangement in Octocorallia. To examine its phylogenetic position of *H. yukibana*, we performed molecular phylogenetic analyses using publicly available mitogenomes of taxa belonging to Clavulariidae, as well as species formerly assigned to this family. We aligned 18,524 nucleotide positions from 14 protein-coding genes and two rRNA genes. *H. yukibana* collected in this study was clustered with three *Clavularia* species (family Clavulariidae).

**Figure 1.**
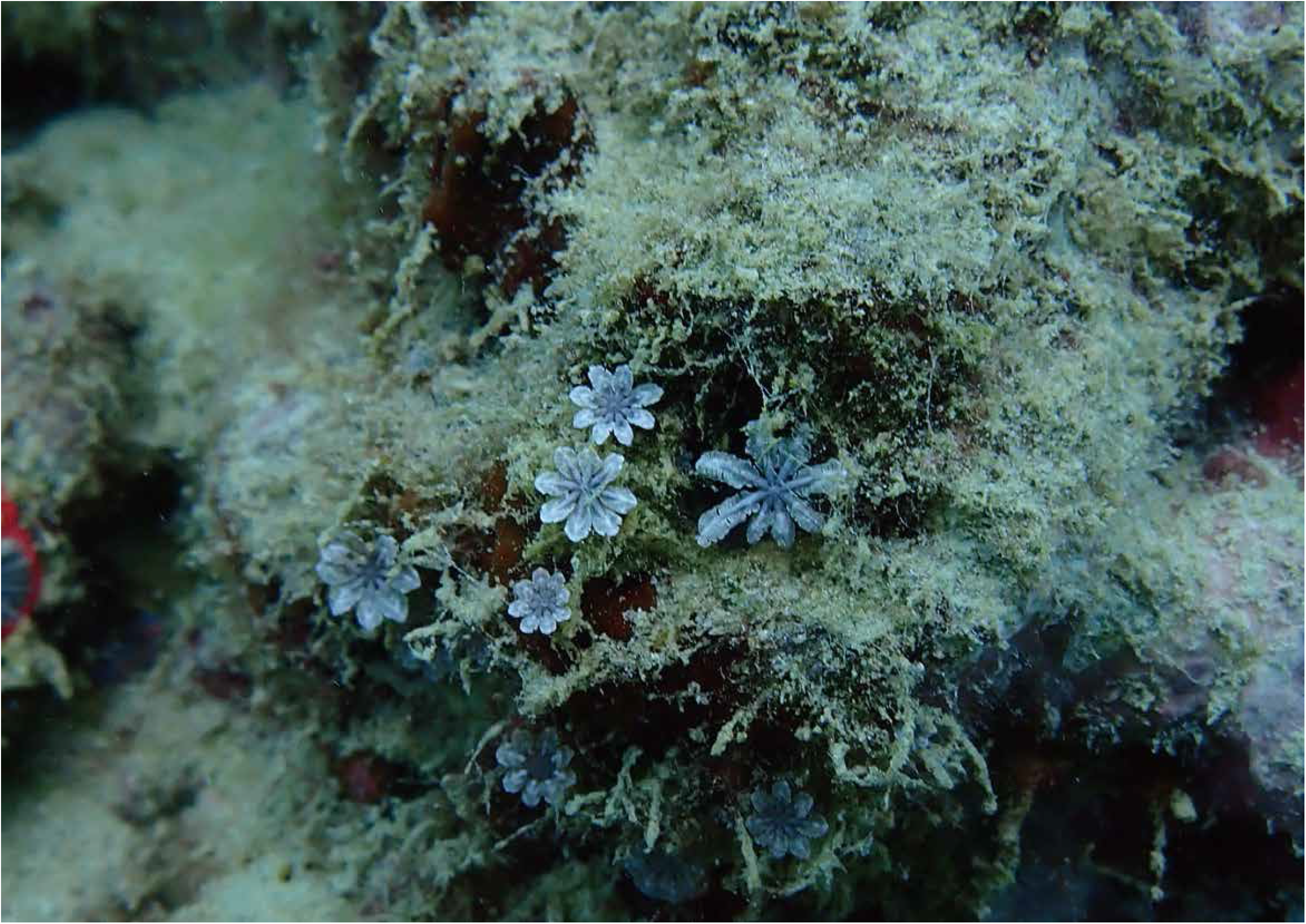
A photograph of *Hanabira yukibana*. Photograph was taken by Tatsuki Koido.

**Figure 2.**
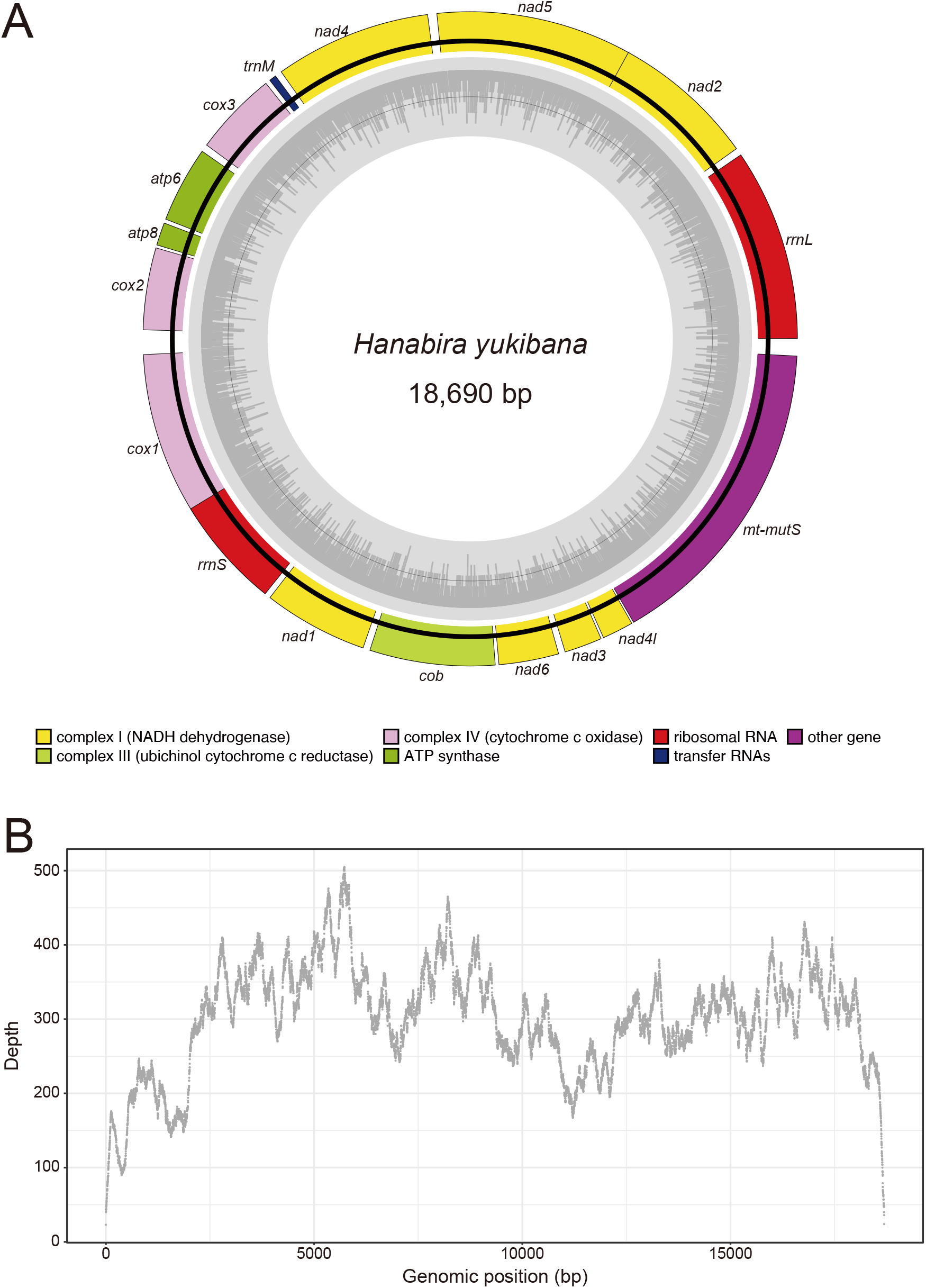
The complete mitochondrial genome of *Hanabira yukibana* and read coverage. **(A)** Inner circles (grey) indicate GC contents. NADH dehydrogenase (yellow), ubichinol cytochrome c reductase (light-green), cytochrome c oxidase (pink), ATP synthase (green), tRNA (blue), rRNAs (red), and other gene (purple). Circular genomes were visualized with OGDRAW. **(B)** Coverage was calculated with BamDeal. The average depth is 279x.

## Discussion and Conclusion

In this study, we reported first mitogenome of *Hanabira yukibana*. Overall, the phylogenetic relationships of *H. yukibana* were congruent with the previous report based on partial mitochondrial gene sets (Lau et al., 2019). It is known that *Clavularia crassa* (NC_069554.1) is clustered with *Paratelesto* sp. (OL616258.1) (family Tubiporidae) (Figure 3), due to the polyphyletic of the genus *Clavularia* reported previously (McFadden et al., 2022; Yoshioka et al., 2025). The complete mitogenome presented here provides an important mitogenomic resource for future taxonomic studies of Clavulariidae and mitogenome evolution among octocorals.

**Figure 3.**
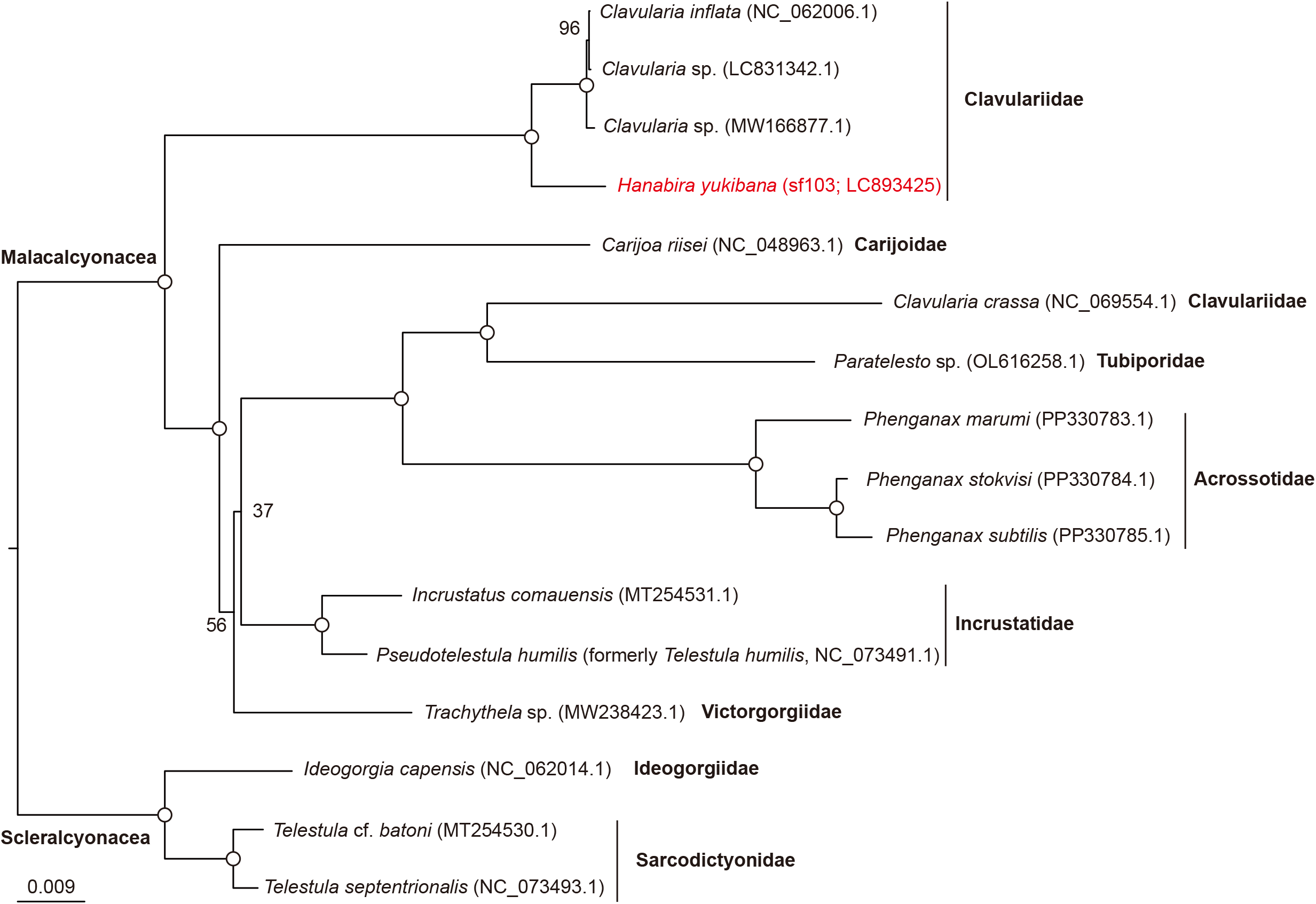
Molecular phylogenetic tree for *Hanabira yukibana* based on mitogenomes. Rooted tree topology was estimated based on 18,524 nucleotide positions comprising 14 protein-coding genes and two rRNA genes. *Hanabira yukibana* is shown in red letter. Accession numbers for mitochondrial genomes are shown in parentheses after scientific names. Open circles indicate 100% bootstrap support (1,000 replicates). The bar indicates expected substitutions per site in aligned regions. The families Ideogorgiidae and Sarcodictyonidae belong to the order Scleralcyonacea, while other families belong to the order Malacalcyonacea. Species used include the following: *Clavularia inflata* (NC_062006.1, Muthye et al., 2022), *Clavularia* sp. (MW166877.1), *Clavularia* sp. (LC831342.1, Yoshioka et al., 2025), *Carijoa riisei* (NC_048963.1, Easton and Hicks, 2020), *Paratelesto* sp. OL616258.1 (Muthye et al.,2022), *Phenganax marumi* (PP330783.1, Poliseno et al., 2025), *Phenganax stokvisi* (PP330784.1, Poliseno et al., 2025), *Phenganax subtilis* (PP330785.1, Poliseno, et al., 2025), *Incrustatus comauensis* (MT254531.1, Poliseno et al., 2021), *Telestula humilis* (NC_073491.1, Poliseno et al., 2021), *Trachythela* sp. (MW238423.1, Zhou et al., 2021), *Ideogorgia capensis* (NC_062014.1, Muthye et al.,2022), *Telestula septentrionalis* (NC_073493.1, Poliseno et al., 2021), and *Telestula* cf. *batoni* (MT254530.1, Poliseno et al., 2021), and *Clavularia crassa* (NC_069554.1).

## Acknowledgments

We thank members of the Sequencing Section at OIST for conducting genome sequencing and members of the Scientific Computing and Data Analysis section at OIST for computing resources.

## Data availability statement

The mitogenome is available in DDBJ/EMBL/GenBank under accession LC893425. The associated BioProject, BioSample and SRA accession numbers are PRJDB17996, SAMD01681849, and DRR747072, respectively.

## CRediT Role

Yuki Yoshioka: Formal analysis; Data curation; Writing – original draft

Tatsuki Koido: Investigation

Megumi Kanai: Funding acquisition; Investigation

Shogo Gishitomi: Investigation

Natsuki Watanabe: Investigation

Noriyuki Satoh: Conceptualization; Funding acquisition; Writing – review & editing

Tomofumi Nagata: Conceptualization; Funding acquisition

## References

Bernt M, Donath A, Jühling F, Externbrink F, Florentz C, Fritzsch G, Pütz J, Middendorf M, Stadler PF. 2013. MITOS: improved de novo metazoan mitochondrial genome annotation. Mol Phylogenet Evol 69, 313–319. Doi: 10.1016/j.ympev.2012.08.023.

Bilewitch JP, Degnan SM. 2011. A unique horizontal gene transfer event has provided the octocoral mitochondrial genome with an active mismatch repair gene that has potential for an unusual self-contained function. BMC Evol Biol 11, 228. Doi: 10.1186/1471-2148-11-228.

Brockman SA, McFadden CS. 2012. The mitochondrial genome of Paraminabea aldersladei (Cnidaria: Anthozoa: Octocorallia) supports intramolecular recombination as the primary mechanism of gene rearrangement in octocoral mitochondrial genomes. Genome Biol Evol 4, 994–1006. Doi: 10.1093/gbe/evs074.

Brugler MR, France SC. 2008. The mitochondrial genome of a deep-sea bamboo coral (Cnidaria, Anthozoa, Octocorallia, Isididae): genome structure and putative origins of replication are not conserved among octocorals. J Mol Evol 67, 125–136. Doi: 10.1007/s00239-008-9116-2.

Cairns SD. 2007. Deep-water corals: an overview with special reference to diversity and distribution of deep-water scleractinian corals. Bull Mar Sci 81, 311–322. URL: http://hdl.handle.net/10088/7536.

Dinesen Z. 1983. Patterns in the distribution of soft corals across the central Great Barrier Reef. Coral Reefs 1, 229–236. Doi: 10.1007/BF00304420.

Easton EE, Hicks D. 2020. Complete mitochondrial genome of Carijoa riisei (Duchassaing & Michelotti, 1860) (Octocorallia: Alcyonacea: Stolonifera: Clavulariidae). Mitochondrial DNA Part B, 5(2), 1826–1827. Doi: 10.1080/23802359.2020.1750998.

Greiner S, Lehwark P, Bock R. 2019. OrganellarGenomeDRAW (OGDRAW) version 1.3. 1: expanded toolkit for the graphical visualization of organellar genomes. Nucleic Acids Res 47:W59–W64. Doi: 10.1093/nar/gkz238.

Hogan RI, Hopkins K, Wheeler AJ, Allcock AL, Yesson C. 2019. Novel diversity in mitochondrial genomes of deep-sea Pennatulacea (Cnidaria: Anthozoa: Octocorallia). Mitochondrial DNA Part A 30, 764–777. Doi: 10.1080/24701394.2019.1634699.

Jin JJ, Yu WB, Yang JB, Song Y, DePamphilis CW, Yi TS, Li DZ. 2020. GetOrganelle: a fast and versatile toolkit for accurate de novo assembly of organelle genomes. Genome Biol 21, 1–31. Doi: 10.1186/s13059-020-02154-5.

Lau YW, Stokvis FR, Imahara Y, Reimer JD. 2019. The stoloniferous octocoral, Hanabira yukibana, gen. nov., sp. nov., of the southern Ryukyus has morphological and symbiont variation. Contrib. Zool 88(1), 54–77. Doi: 10.1163/18759866-20191355.

Martin M. 2011. Cutadapt removes adapter sequences from high-throughput sequencing reads. EMBnet. journal 17, 10–12. Doi: 10.14806/ej.17.1.200.

McFadden CS, Sánchez JA, France SC. 2010. Molecular phylogenetic insights into the evolution of Octocorallia: a review. Integr Comp Biol 50, 389–410. Doi: 10.1093/icb/icq056.

McFadden CS, Van Ofwegen LP, Quattrini AM. 2022. Revisionary systematics of Octocorallia (Cnidaria: Anthozoa) guided by phylogenomics. Bull Soc Syst Biol 1(3). Doi: 10.18061/bssb.v1i3.8735.

Muthye V, Mackereth CD, Stewart JB, Lavrov DV. 2022. Large dataset of octocoral mitochondrial genomes provides new insights into mt-mutS evolution and function. DNA repair 110, 103273. Doi: 10.1016/j.dnarep.2022.103273.

Pante E, Saucier EH, France SC. 2013. Molecular and morphological data support reclassification of the octocoral genus Isidoides. Invertebr Syst 27, 365–378. Doi: 10.1071/IS12053.

Park E, Hwang DS, Lee JS, Song JI, Seo TK, Won YJ. 2012. Estimation of divergence times in cnidarian evolution based on mitochondrial protein-coding genes and the fossil record. Mol Phylogenet Evol 62, 329–345. Doi: 10.1016/j.ympev.2011.10.008.

Pont-Kingdon GA, Okada NA, Macfarlane JL, Beagley CT, Wolstenholme DR, Cavalier-Smith T, Clark-Walker GD. 1995. A coral mitochondrial mutS gene. Nature 375, 109–111. Doi: 10.1038/375109b0.

Poliseno A, Altuna A, Puetz LC, Mak SS, Ríos P, Petroni E, McFadden CS, Sørensen MV, Gilbert MTP. 2021. An integrated morphological–molecular approach reveals new insights on the systematics of the octocoral Telestula humilis (Thomson, 1927) (Octocorallia: Alcyonacea: Clavulariidae). Invertebr Syst 35(3), 261–281. Doi: 10.1071/IS20009.

Poliseno A, Quattrini AM, Lau YW, Pirro S, Reimer JD, McFadden CS. 2025. New mitochondrial gene order arrangements and evolutionary implications in the class Octocorallia. Mitochondrial DNA Part A 35(1-2), 23–33. Doi: 10.1080/24701394.2024.2416173.

Uda K, Komeda Y, Koyama H, Koga K, Fujita T, Iwasaki N, Suzuki T. 2011. Complete mitochondrial genomes of two Japanese precious corals, Paracorallium japonicum and Corallium konojoi (Cnidaria, Octocorallia, Coralliidae): notable differences in gene arrangement. Gene 476, 27–37. Doi: 10.1016/j.gene.2011.01.019.

Williams GC, Cairns SD. 2019. Systematic list of valid octocoral genera. Accessed through: https://researcharchive.calacademy.org/research/izg/OCTOCLASS.htm on 2025-09-08.

Yoshioka Y, Kanai M, Gishitomi S, Arakaki N, Koido T, Shinzato C, Inoue J, Nagata T, Satoh N. 2025. Extensive mitochondrial genomic analyses reveal dynamic gene order rearrangements in the class Octocorallia (Cnidaria: Anthozoa). Gene Rep 38, 102111. Doi: 10.1016/j.genrep.2024.102111.

Zhou Y, Feng C, Pu Y, Liu J, Liu R, Zhang H. 2021. The first draft genome of a cold-water coral Trachythela sp. (Alcyonacea: Stolonifera: Clavulariidae). Genome Biol Evol 13(2), evaa265. Doi: 10.1093/gbe/evaa265.

